# Neuron type-specific mechanical regulation of voltage-gated Ca^2+^ channels and excitability in hippocampal and trigeminal ganglion neurons

**DOI:** 10.1101/123364

**Authors:** Sisi liu, Yang Yu, Xiaoan Wu, Liying Huang, Yuchen Qi, Lin-Hua Jiang, Hucheng Zhao

## Abstract

Increasing evidence suggests that the mechanical properties of extracellular matrix regulate central and peripheral neuronal functions. We thus investigated the Ca_V_ channels in hippocampal and trigeminal ganglion (TG) neurons cultured on substrates with different stiffness. Patch-clamp current recordings showed that stiff substrate augmented the Ca_V_ channel currents in hippocampal and TG neurons and additionally induced a leftward shift in the voltage-dependent channel activation curve in small TG neurons. Combination with using selective channel blockers revealed that substrate stiffness preferentially regulated the N-type channel current in hippocampal and medium TG neurons but the T-type channel current in small TG neurons. Current-clamp recordings further demonstrated that stiff substrate enhanced the excitability of small TG neurons, which was ablated by blocking the T-type channel. Treatment of neurons on the stiff substrate with low-dose blebbistatin reduced both the N-type channel current in hippocampal and medium TG neurons and the T-type channel current in small TG neurons to the levels in neurons on the soft substrate, whereas treatment of neurons on the soft substrate with calcium A increased both the N-type channel current in hippocampal and medium TG neurons and the T-type channel current in small TG neurons to the levels in neurons on the stiff substrate, thus consistently supporting critical involvement of actomyosin in mechanical sensing. Taken together, our results reveal neuron type-specific mechanical regulation of the Cav channels and excitability in the nervous system. Such information is useful for neural tissue engineering and regeneration.

## Introduction

Many cells types in multi-cell organisms are able to detect and respond to extracellular chemical signals and also sense physical clues in their surroundings (Schiffhauer et al., 2016). Increasing evidence suggests that central and peripheral neurons are responsive to the mechanical properties of extracellular matrix (ECM) and cell-supporting substrates with profound change in their morphological patterning, neurite outgrowth and process branching (Georges et al., 2006; Kostic et al., 2007; Previtera et al., 2010; Sur et al., 2013; Previtera et al., 2013; Fabbro et al., 2012; Cellot et al., 2011; Tang et al., 2013; Koch et al., 2012). Thus, substrates with stiffness levels similar to the *in vivo* microenvironments tend to promote neuronal growth over glia growth in the cortical cultures (Georges et al., 2006). Stiff substrates stimulate dendrite arborization of hippocampal neurons, whereas soft substrates result in weak adhesion and facilitate neurite extension and/or retraction and thereby increase the frequency of extension-retraction events, leading to neurite motility (Previtera et al., 2010). In agreement with these findings, emerging evidence shows that substrate stiffness regulates neuronal network formation and activity (Lantoine et al., 2016; Zhang et al., 2014). However, the mechanical properties of the *in vivo* surroundings for the central and peripheral neurons are noticeably different. Brain tissues contain a significant number of glial cells and represent one of the softest tissues in the body (Lu et al., 2006), whereas peripheral neurons experience different mechanical microenvironments when they navigate through various organs and tissues (Franze and Guck, 2010). There is evidence that central and peripheral neurons can respond differently to mechanical signals. For instance, dorsal root ganglion neurons (DRG) exhibit strong neurite outgrowth on soft substrates and noticeably less growth on stiff substrates, but hippocampal neurite outgrowth is independent of substrate stiffness (Georges et al., 2006; Koch et al., 2012). It has been proposed that many cell types detect the mechanical properties by applying traction forces through the actomyosin stress fibers to the substrates at focal adhesions (Lee and Kumar, 2016; Kobayashi and Sokabe, 2010). It is poorly understood whether neuron-specific sensitivity or regulation of substrate stiffness arise from difference in the mechanosensing mechanisms and the downstream signaling pathways in the central and peripheral neurons.

The voltage-gated Ca^2+^ (Cav) channels in neurons are important in transducing electrical signals to intracellular Ca^2+^ signaling that has a role in regulating a wide range of functional processes such as neurotransmission and synaptic plasticity in the nervous system (Catterall and Swanson, 2015; Proft and Weiss, 2015; Zamponi, 2016; Cheong and Shin, 2013). We have recently reported substrate stiffness regulates the Cav channels in hippocampal neurons with strong consequences on the neuronal network formation and activity (Zhang et al., 2014). It is well documented that both central and peripheral neurons express multiple Cav channels (Catterall and Swanson, 2015; Proft and Weiss, 2015; Zamponi, 2016), but the identity of the Cav channels regulated by the mechanical properties remains unknown. It is also unknown whether and which Cav channels in the peripheral neurons are under mechanical regulation. Therefore, in this study we used hippocampal neurons and trigeminal ganglion (TG) neurons to investigate whether substrate stiffness imposes similar regulation of neuronal Cav channels in the central and peripheral neurons and targets the same type of Cav channels as well as to examine whether actomyosin-mediated mechanosensing mechanism is critically engaged in mechanical regulation of neuronal Cav channels. Our study provides the first evidence to show that substrate stiffness regulation of Cav channels and excitability is neuron type-specific and actomyosin-mediated mechanosensing mechanism is critical in mechanical regulation of different Cav channels in the central and peripheral neurons. These findings offer molecular insights into mechanical regulation of neuronal cell functions and are useful for development of novel scaffolds for neural tissue engineering or regeneration.

## Materials and Methods

### Preparation of PDMS substrates

PDMS were prepared according to published procedures (Zhang et al., 2014; Beaudoin et al., 2012). Briefly, curing agent (sylgard184; Dow corning Corp) were mixed with base agent in a mass ratio of 10:1 or 50:1 and coated with laminin used to culture hippocampal and TG neurons. The Young’s moduli were 457 ± 39 and 46 ± 11 kPa in 1:10 and 1:50 PDMS, respectively, measured by the spherical indentation method (Zhang et al., 2014), which are referred to as stiff and soft substrates in this study.

### Hippocampal and TG neuron culture preparation

All the procedures of using animals were in accordance with the guidelines approved by Tsinghua University. Mouse hippocampal neurons were isolated from one day old pups as previously described (Zhang et al., 2014). In brief, after the blood vessels and meninges were removed, hippocampal tissues were collected into tubes in cold Hank’s balanced salt solution (HBSS), and rinsed twice with HBSS solutions. Tissues were digested in TrypIE (Sigma) for 15 min at 37°C and gently triturated using pipette tips to cell suspension in DMEM medium supplemented with 10% fetal bovine serum (FBS) and 2% B-27 (Life Technologies). Cells were maintained at 37°C in humidified atmosphere with 5% CO_2_. Arabinofuranosyl cysteine at 1 mg/ml was added to culture medium 2-3 days after initial seeding to inhibit glial proliferation. TG tissues were collected from 1-2 day old mice as previously described (Tao et al., 2012). In brief, tissues treated with 5 mg/ml trypsin for 15 min. The culture medium contained 5% FBS, 10 ng/ml nerve growth factor and 10 ng/ml glial cell line-derived neurotrophic factor, and was replaced every 3 days.

### Immunocytochemistry and image analysis

Cells were washed with PBS pre-warmed at 37°C, fixed with 4% paraformaldehyde in PBS for 10 min, permeabilized in 0.1% Triton X-100 PBS for 5 min, and washed with PBS four times, each time for 15 min. For MAP2 straining, cells were blocked with 5% bovine serum albumin in PBS for 60 min, and incubated with primary rabbit polyclonal MAP2 antibody (Abcam) at a dilution of 1:500 overnight. After extensive wash with PBS, cells were incubated with secondary Alexa Fluor568 conjugated goat anti-rabbit IgG antibody (Molecular Probes) at 1:1000 for 60 min.

### Size distribution and IB4 labelling of TG neurons

Differential interference contrast (DIC) images of cultured TG neurons were captured using a Nikon TE2000S inverted microscope equipped with a CoolSnapEZ camera and analyzed with SimplePCI software (Hamamatsu). Soma diameter was converted from cross-sectional area measured from the DIC image. TG neurons were incubated with Alexa Fluor 594-conjugated IB4 (3 μg/ml) for 10 min. The Alexa Fluor 594 fluorescence on soma membrane was detected after washing for 10 min.

### Ba^2+^ current recording

TG and hippocampal neurons 4-6 DIV were recorded with 5 mM Ba^2+^ as charge carrier. Whole cell patch-clamp recordings were performed at room temperature using a MultiClamp 700B amplifier and pClamp 10 (Molecular Devices). The recording chamber was perfused with extracellular solution (0.5 ml/min) containing (in mM): 20 CsCl, 140 TEA-Cl, 5 BaCl**2**, 10 HEPES, 25 glucose, pH 7.3 with TEA-OH, and 310 mosmol/kg H_2_O. The pipette solution contained (in mM): 110 CsCl, 10 EGTA, 4 ATP-Mg, 0.3 GTP-Na, 25 HEPES, 10 Tris-phosphocreatine, 20 U/ml creatine phosphokinase, pH 7.3 with CsOH, and 290 mosmol/kg H_2_O. Recording pipettes had <3.5 ΜΩ resistance (average 2.7 ΜΩ). Series resistance (<15 ΜΩ, average 10.0 ΜΩ) was compensated by 80%. The cell capacitance and series resistance were constantly monitored during recording. Current traces were corrected with online P/6 trace subtraction. Signals were filtered at 1 kHz and digitized at 10 kHz except for the tail currents, which were filtered at 10 kHz and digitized at 100 kHz. The DIC images of neurons were captured before recording to estimate the soma size of TG neurons.

To determine the current-voltage (I-V) relationships curves of high-voltage-activated (HVA) Ca^2+^ channels, neurons were held at −80 mV and were depolarized from −80 to +100 mV with 10 mV increments for 40 ms and returned to −80 mV. The voltage dependence of Cav channel activation was determined by the tail currents which were normalized to the largest current and the activation curve fitted by a Boltzmann equation. To dissect whole-cell Ba^2+^ currents through different HVA types of Cav channels, a paradigm described in a previous study was used (Zhang et al., 2002; Tao et al., 2012). The Ba^2+^ currents were evoked by ramp depolarization pulses from −80 to +100 mV at 1.8 mV/ms every 10 s. L-, N-, and P/Q-type Cav channel currents were defined by the current components sensitive to 10 μM nifedipine, 2 μM ω-conotoxin-GVIA (ω-CTx-GVIA) and 0.5 μM ω-agatoxin-IVA (ω-Aga-IVA), respectively. The residual current that was inhibited by 100 μM Cd^2+^ was regarded to be medicated by the R-type Cav channel. The peak current was normalized by the membrane capacitance to derive the current density.

The following protocol described in a previous study based on that the majority of T-type channels are inactivated at more depolarizing holding potentials (Zhang et al., 2002) was used to dissect the low-voltage-activated (LVA) T-type Cav channel currents. Most, if not all, HVA channel currents were blocked by bath-applying a cocktail of channel blockers, comprising 10 μM nifedipine, 2 μM ω-CTx-GVIA, 0.5 μM ω-Aga-IVA and 0.2 μM SNX482 (Zhang et al., 2002). The Ba^2+^ currents evoked by 40 ms depolarization to −40 mV from a holding potential (V_h_) of −60 or −110 mV, and the T-type channel current was derived by digitally subtracting the current evoked at the V_h_ of −60 mV from that measured at the Vh of −110 mV. This eliminated the small but substantial residual HVA channel currents that was not blocked by the channel blocker cocktail (Cellot et al., 2011). Voltage dependence of T-type Cav channel activation in TG neurons was also determined from the tail currents. The HVA Cav channel currents were minimized with the channel blocker cocktail. Neurons held at either −60 or −110 mV were depolarized from −80 to 30 mV with 10-mV increments for 40 ms and then repolarized back to the V_h_. The amplitude of tail currents, derived by digital subtraction as described above, were normalized to the largest current and were fitted by a Boltzmann equation.

### Current-clamp recording

The recording chamber was perfused with Tyrode solution (0.5 ml/min) containing (in mM): 130 NaCl, 2 KCl, 2 CaCl_2_, 2 MgCl_2_, 25 HEPES, 30 glucose, pH 7.3 with NaOH, and 310 mosmol/kg H_2_O. The pipette solution contained (in mM): 130 K-gluconate, 7 KCl, 2 NaCl, 1 MgCl_2_, 0.4 EGTA, 4 ATP-Mg, 0.3 GTP-Na, 10 HEPES, 10 Tris-phosphocreatine, 20 U/ml creatine phosphokinase, pH 7.3 with KOH, and 290 mosmol/kg H_2_O. Recording pipettes had resistance of <4.5 MΩ. Series resistance (<20 MΩ) was not compensated. Signals were filtered at 10 kHz and digitized at 50 kHz. After whole-cell access was established, the amplifier was switched to the current-clamp mode to measure the resting membrane potential (V_rest_). The input resistance (R_in_) was calculated by measuring the change of membrane potential in response to a 20-pA hyperpolarizing current injection from V_rest_. Neurons were excluded from analysis if the V_rest_ was higher than −40 mV or R_in_ was < 200 MΩ. To test neuronal excitability, neurons were held at V_rest_ and were injected with 1-s depolarizing currents in 25-pA incremental steps until at least one action potential (AP)-like spike was elicited. The rheobase was defined as the minimum amount of current to elicit ≥1 AP. The first AP elicited using this paradigm was used to measure the AP threshold (membrane potential at which dV/dt exceeds 10 V/s), amplitude and half width. The amplitude of after hyperpolarization (AHP) was measured from the single AP elicited by injecting 0.5∼2-nA depolarizing current for 0.5 ms. To measure the spike frequency in response to suprathreshold stimuli, neurons were injected with 1-s depolarizing currents at one-, two-, and threefold rheobase.

### Drug applications

Nifedipine (Sigma-Aldrich) was dissolved in 100% ethanol to generate 10 mM stock solution. All peptide Cav channel blockers were purchased from Peptides International (Louisville, KY), reconstituted in 1 mg/ml cytochrome *c* (Sigma-Aldrich) at 100 times concentrations, and stored at −80°C in aliquots. Before the application of Cav channel blockers, the perfusion in the recording chamber was stopped and cytochrome *c* was added to the bath at the final concentration of 0.1 mg/ml. Subsequently, the blockers were applied sequentially and cumulatively to the recording chamber. Blebbistatin (Sigma) and caliculin A (Sigma) were added in the culture medium for 24 h with a final concentration of 3 μM and 20 nM, respectively, before cells were used.

### Data presentation and statistical analysis

Data are reported as means ± standard error of mean (SEM). The neuron density was derived from MAP2-positive cells that were counted in fluorescent images and converted to the number of neurons per mm^2^, and the soma area of individual MAP2-positive cells was estimated using ImageJ. Electrophysiological data were analyzed with Clampfit (Molecular Devices) and Origin (OriginLab) softwares. The voltage-dependent activation curves were fitted to the Boltzmann equation: I/I_max_ = 1/[1+exp[(V-V_1/2_)/k]), where I_max_, V_1/2_ and k represents the maximal current, half maximal current activation voltage and slope, respectively. Statistical significance between groups was assessed by two-tail *t*-test or ANOVA with post hoc Bonferroni test. Statistical significance between the I-V curves and between the spike frequency curves was assessed by repeated-measures ANOVA (RM ANOVA) with post hoc Bonferroni test.

## Results

### Substrate stiffness has no effect on intrinsic properties of hippocampal and TG neurons

Hippocampal and TG neurons cultured on the soft and stiff substrates over a period of 2-3 days showed typical cell body size, morphology, neurite growth and extension (Fig. S1A). Confocal imaging analysis of MAP2-positive cells revealed no difference in hippocampal and TG neuron density cultured on the stiff and soft substrates (Fig. S1B) as reported in our previous study [14]. TG neurons are heterogeneous and can are divided, according to the soma size, into small (< 25 μm), medium (25-35 μm) and large TG neurons (> 35 μm) subpopulations, and the small TG neurons can be further subdivided into IB4^+^and IB4^−^ groups (Tao et al., 2012). There were very few large TG neurons in our preparations. Culturing on substrates with a 10-fold difference in stiffness had no effect on TG neuron subpopulations with 23%, 28% and 49% for medium, IB4+ and IB4^−^ neurons on the soft substrate, and 22%, 27% and 51% on the stiff substrate (Tab. 1). We also characterized the intrinsic electrical properties of hippocampal, medium and small TG neurons, including capacitance, input resistance, resting membrane potential, and AP amplitude. All these properties were virtually the same for all types of neurons cultured on the soft and stiff substrates (Tab. 2 and Tab. 3).

**Table 1.** Subpopulations of TG neurons cultured on the soft and stiff substrates.

| Neuron type | % of total neurons (soft) | % of total neurons (stiff) |
| --- | --- | --- |
| Small IB4<sup>-</sup> | 49 | 51 |
| Small IB4<sup>+</sup> | 28 | 27 |
| Medium sized | 23 | 22 |

n = 450 neurons in 6 preparations examined for each case.

**Table 2.** Electrophysiological properties of hippocampal and medium TG neurons on the stiff and soft substrates.

| Parameter | Hippocampal neurons |  | Medium size TG neurons |  |
| --- | --- | --- | --- | --- |
|  | Stiff | soft | stiff | soft |
| Diameter ($\mu\text{m}$) | $15.3 \pm 1.2$ | $15.9 \pm 1.6$ | $28.3 \pm 2.1$ | $27.8 \pm 1.8$ |
| Capacitance (pF) | $33.3 \pm 2.1$ | $31.4 \pm 2.6$ | $26.7 \pm 1.8$ | $25.5 \pm 2.5$ |
| $R_{\text{in}}$ (MΩ) | $1226 \pm 44$ | $1198 \pm 67$ | $1099 \pm 72$ | $1091 \pm 68$ |
| $V_{\text{rest}}$ (mV) | $-60 \pm 4$ | $-61 \pm 5$ | $-56 \pm 6$ | $-54 \pm 4$ |
| Amplitude (AP) | $106 \pm 8$ | $102 \pm 9$ | $104 \pm 6$ | $103 \pm 6$ |

n = 11-12 neurons examined for each parameter under each condition. No significant difference in electrophysiological properties between neurons cultured on soft and stiff substrates.

**Table 3.**
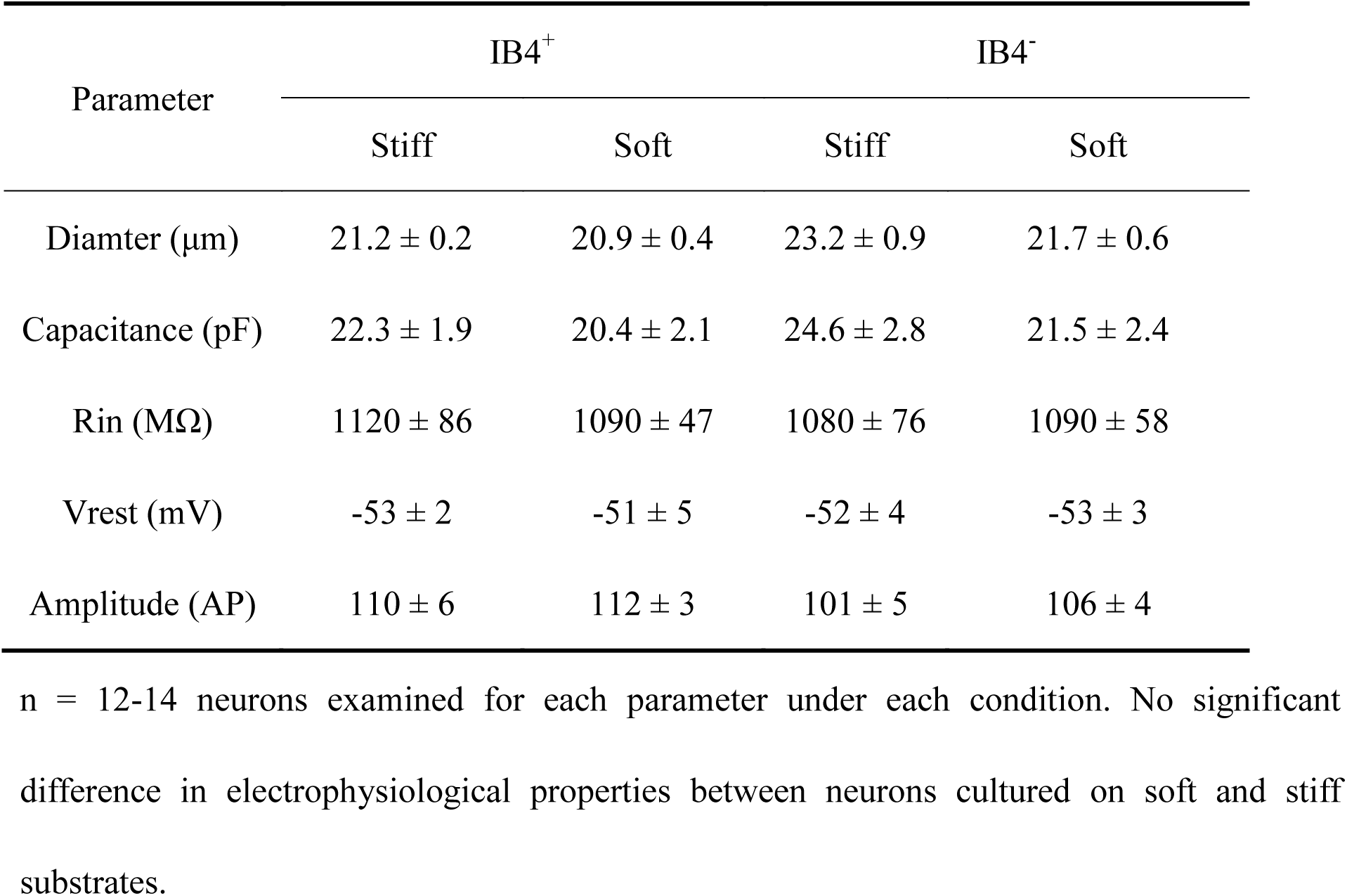
Electrophysiological properties of small TG neurons on stiff and soft substrates.

### Stiff substrate upregulates total Ca_V_ channel currents in hippocampal and TG neurons

We performed voltage-clamp whole-cell current recording to compare the effects of substrate stiffness on the Ca_V_ channel currents in hippocampal neurons, and medium, small IB4^+^ and IB4^−^ TG neurons. Membrane depolarization evoked significant Ca_v_ channel currents in all these neuron types cultured on both soft and stiff substrates (Fig. 1A-D), with the typical bell-shaped I-V relationship curve (Fig. 1E-H). The inward currents began to appear at approximately −40∼−50 mV, peaked at −20∼−10 mV and reversed at +50 mV (Fig. 1E-H). The peak inward current amplitude was significantly greater in hippocampal, small and medium TG neurons on the stiff substrate than on the soft substrate (Fig 1 E-H). Interestingly, there is noticeable difference in the extent, to which stiff substrate increased the Cav channel currents in different types of neurons, in the order from high to low: hippocampal neuron > medium TG neuron > small IB4^+^ TG neuron ∼ small IB4^−^ neuron (Fig. 1I). Taken together, these results show that substrate stiffness regulates the Cav channels in the central and peripheral neurons but there is difference in such regulation.

**Fig. 1.**
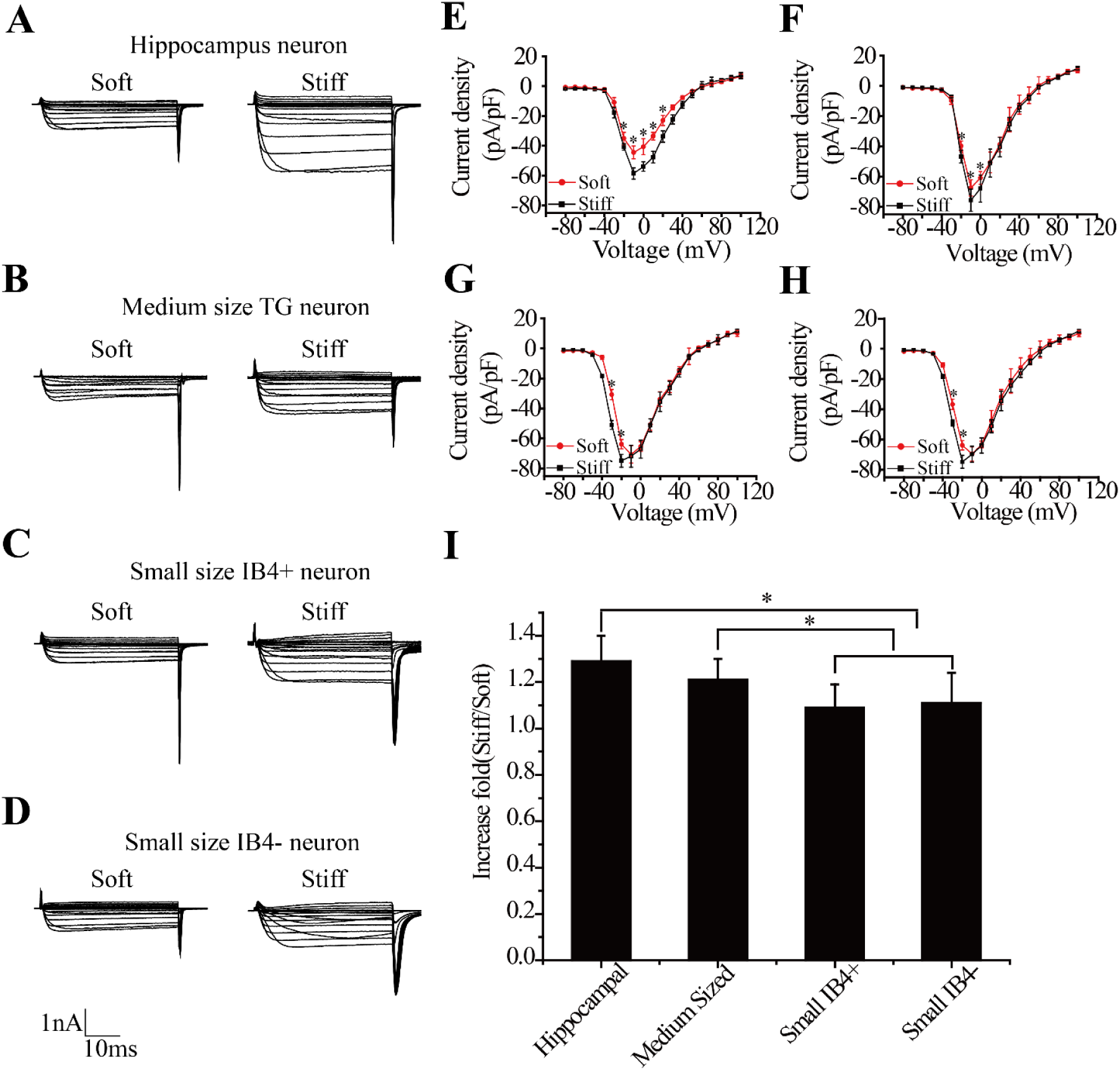
Stiff substrate enhances Cav channel currents in hippocampal and TG neurons. **A-D.** Representative patch-clamp recording of whole-cell Cav channel currents using Ba^2+^ as charge carrier from indicated neurons on the soft and stiff substrates. **E-H**. Summary of the I-V curves of Cav channel currents recorded from indicated neurons on soft and stiff substrates, with the number of neurons examined as follows: 10 hippocampal neurons (E), 11 medium TG neurons (F), 11 small IB4^+^ TG neurons (G), and 11 small IB4^−^ TG neurons (H) from three preparations. **I**. Summary of the mean increase in the peak Cav channel current density by stiff substrate relative to soft substrate. *, p < 0.05 denoting significant difference between indicated groups.

We next examined the effects of substrate stiffness on voltage dependence of the Cav channel activation. The voltage-dependent Cav channel activation curves, determined based on the tail currents, were virtually superimposed for hippocampal and medium TG neurons on the soft and stiff substrates (Fig. 2A-B). This is further indicated by no significant difference in the V_1/2_ values derived by fitting the activation curves. The V_1/2_ values were −6.8 ± 0.7 mV and −5.3 ± 0.6 mV for hippocampal neurons on the stiff and soft substrates respectively (Fig. 2A), and the V_1/2_ values were −7.8 ± 0.8 mV and −6.9 ± 0.7 mV for medium TG neurons on the stiff and soft substrates, respectively (Fig. 2B). In striking contrast, the voltage-dependent activation curves in both small IB4^+^ and IB4^−^ TG neurons on the stiff substrate showed a remarkable leftward shift compared to those in neurons on the soft substrate (Fig. 2C-D). The V_1/2_ values were −24.4 ± 2.1 mV and −7.4 ± 1.1 mV for IB4^+^1 neurons on the stiff and soft substrates (Fig. 2C; p < 0.05), respectively, and −25.3 ± 1.7 mV and −7.9 ± 0.9 mV for IB4^−^ neurons on the stiff and soft substrates, respectively (Fig. 2D; p < 0.05). These results provide strong evidence to indicate neuron type-specific mechanical regulation of the Cav channels.

**Fig. 2.**
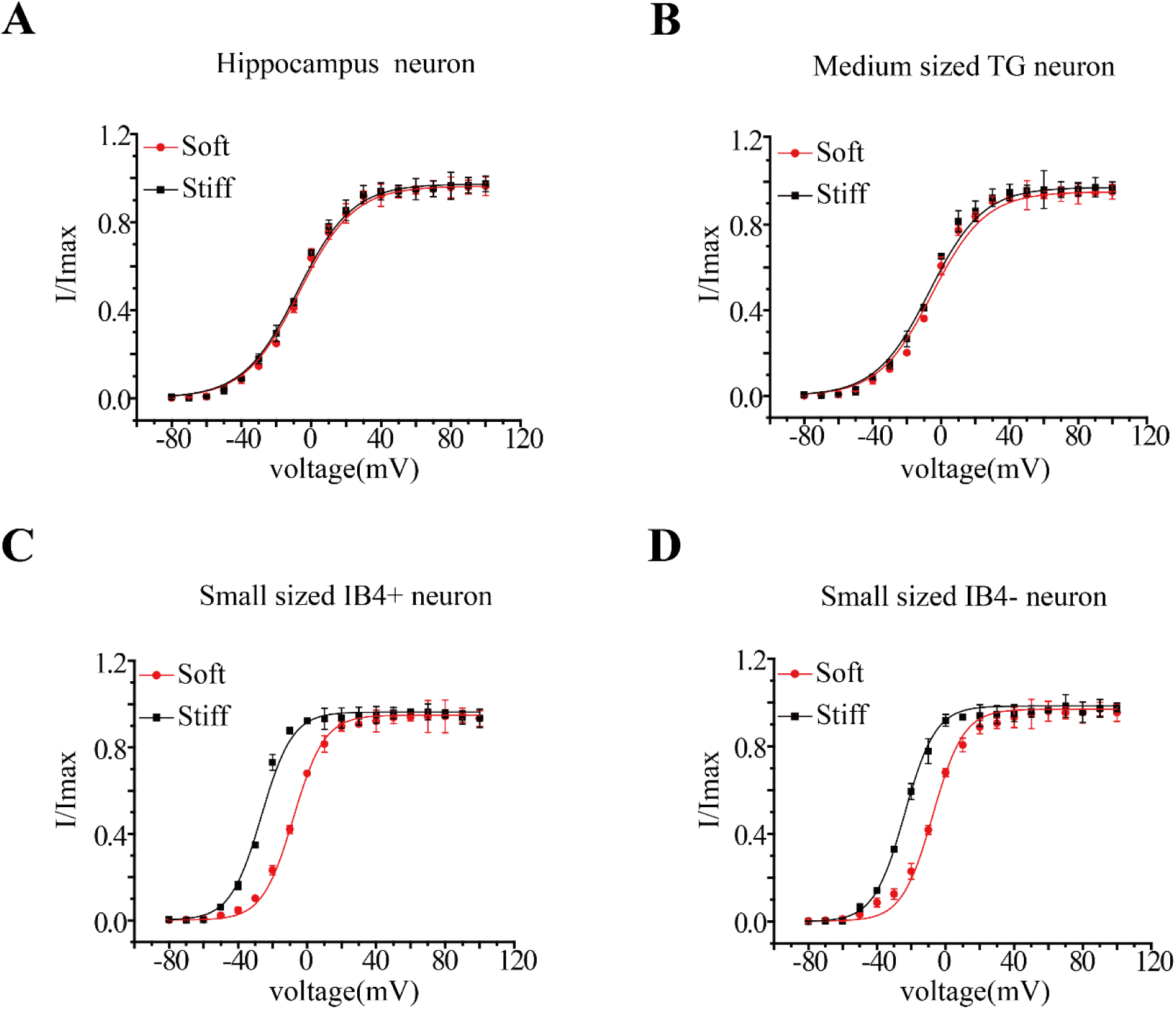
Stiff substrate induces a left-shift in voltage-dependent Cav channel activation in small TG neurons but no effect in hippocampal and medium TG neurons. **A-D.** Summary of the voltage-dependent Cav channel activation in indicated neurons on soft and stiff substrates from three preparations, determined by measuring tail currents. The number of neurons examined is as follows: 10 hippocampal neurons (A), 11 medium sized TG neurons (B), 11 small IB4^+^ TG neurons (C), and 11 small IB4^−^ TG neurons (D).

### Stiff substrate augments N-type channel currents in hippocampal and medium TG neurons, but not in small TG neurons

Hippocampal and TG neurons are known to express multiple HVA L-, N-, P/Q- and R-types, and LVA T-type Cav channels (Tao et al., 2012; Cao et al., 2004). To examine whether the HVA Cav channels and which of them are regulated by substrate stiffness, we measured the Cav channel current components through each of the HVA Ca^2+^ channels with a holding potential (V_h_) of −80 mV (to inactivate the LVA T-type channel) and adopting a pharmacological protocol described in previous studies (Tao et al., 2012; Zhang et al., 2002), namely, 2 μM ω-CTx-GVIA, 0.5 μM ω-Aga-IVA, 10 μM nifedipine and 100 μM Cd^2+^ were stepwise added during the recording, to block the N-, P/Q-, L-type and R-type channels, respectively (Fig. 3A). In hippocampal and medium TG neurons, the N-type channel current was significantly greater in neurons on the stiff substrate than on the soft substrate, but there was no difference in the currents mediated by the other HVA Cav channel types (Fig. 3B-C). However, the Ca^2+^ currents mediated by all these HVA Cav channels remained the same between small IB4^+^ and IB4^−^ TG neurons on the soft and stiff substrates (Fig. 3D-E). Characterization of the ω-CTx-GVIA-sensitive currents in hippocampal and medium TG neurons further indicates stiff substrate increased the N-type channel current (Fig. 3F-G, I-J) with no alteration in the voltage-dependent channel activation curve (Fig. 3H and K). Collectively, these results show that substrate stiffness preferentially alters the N-type channel in hippocampal and medium TG neurons, but not in small TG neurons.

**Fig. 3.**
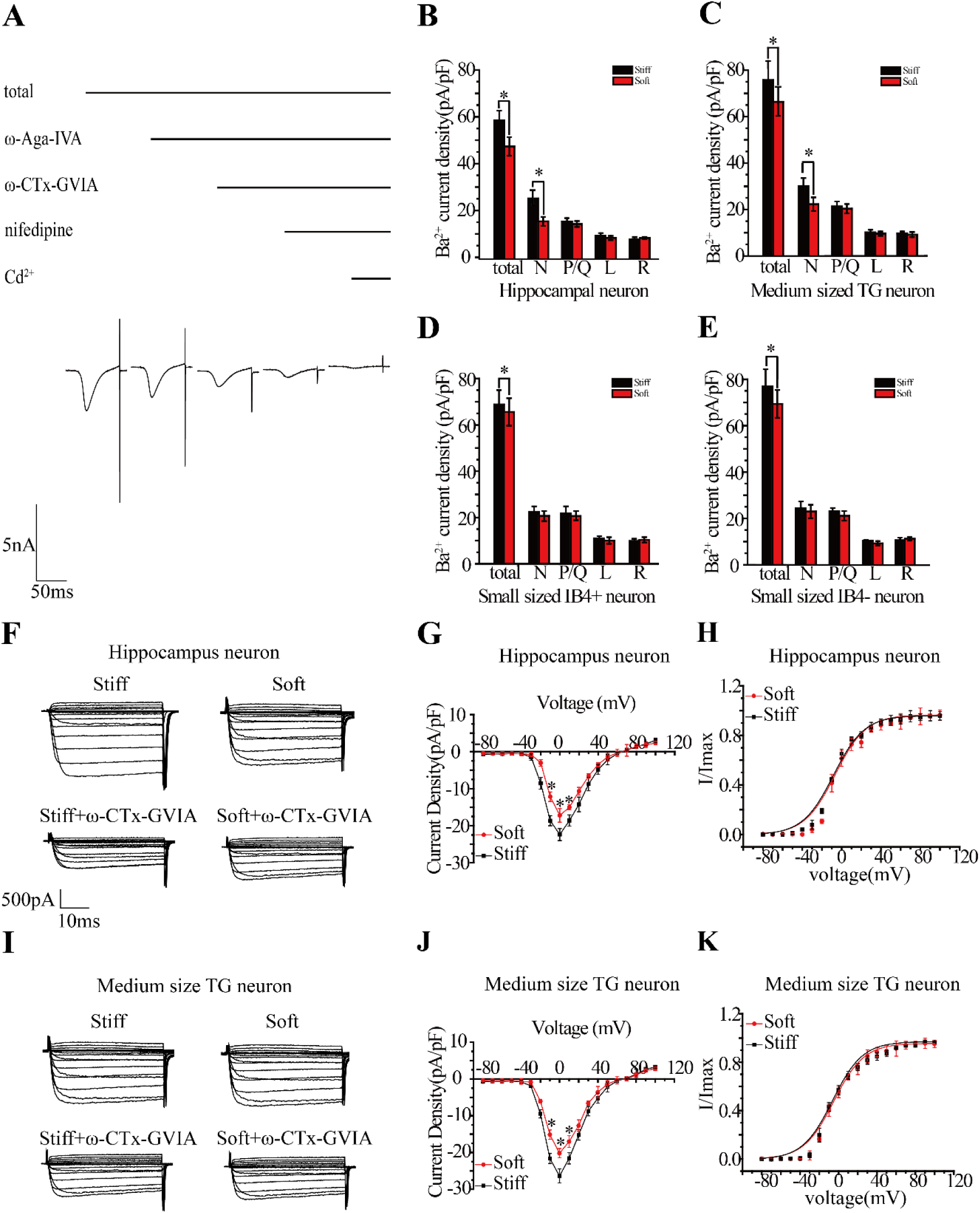
Stiff substrate augments N-type channel currents in hippocampal and medium TG neurons, but not in small TG neurons. **A.** Representative patch-clamp recording of whole-cell Cav channel currents using Ba^2+^ as charge carrier in response to sequential addition of Cav channel type specific blockers (0.5 μM ω-Aga-IVA, 2 pM ω-conotoxin-GVIA, 10 μM nifedipine, and 100 μM Cd^2+^). **B-E**. Summary of the mean peak current densities, showing total HVA Cav channel currents and also currents mediated by different HVA Cav channel types in indicated neurons on the soft and stiff substrates from three preparations, with the number of neurons examined and the membrane potential as follows: 8-11 hippocampal neurons at −10 mV (B), 9-11 medium TG neurons at −10 mV (C), 8-11 small IB4^+^ TG neurons at −20 mV (D), and 9-11 small IB4^−^ TG neuron at −20 mV (E). **F**. Representative recording of whole-cell Cav channel currents in hippocampal neurons on the soft and stiff substrates in the absence (top panels) or presence of 2 μM ω-conotoxin-GVIA (bottom panels). **G-H**. Summary of the I-V curve of peak N-type channel currents (G), and the voltage dependent N-type channel activation determined by measuring tail currents (H), from 10 hippocampal neurons cultured on soft and stiff substrates in three preparations. **I**. Representative recording of whole-cell Cav channel currents in medium TG neurons on the soft and stiff substrates in the absence (top panels) or presence of 2 μM ω-conotoxin-GVIA (bottom panel). **J-K**. Summary of the mean I-V curves of peak N-type channel (J), and the voltage dependent N-type channel activation determined by measuring tail currents (K), from 11 medium TG neurons on soft and stiff substrates in three preparations. *, p < 0.05 denoting significant difference between indicated groups.

### Stiff substrate up-regulates T-type channel currents in small TG neurons, but not in hippcampal and medium TG neurons

Our results above showed that stiff substrate induced a strong leftward shift in the I-V relationship curve as well as an increase in the total Ca^2+^ channel currents in small TG neurons (Fig. 1G-H). We hypothesized these effects were mediated by the LVA T-type Cav channel. To test this hypothesis, we analysed the T-type channel currents in small TG neurons cultured on the stiff and soft substrates, and also in hippocampal and medium TG neurons for direct comparison. The T-type channel currents were isolated by substrating the Ca^2+^ currents induced by depolarizing to −40 mV from the V_h_ of −60 mV (full T-type channel inactivation) from the currents from the V_h_ of −110 mV (no T-type channel inactivation) in the same neurons that were treated prior and during the current recording with a cocktail of HVA Cav channel blockers (10 μM nifedipine, 2 μM ω-CTx-GVIA, 0.5 μM ω-Aga-IVA and 0.2 μM SNX482) (Fig. S2). For hippcampal and medium TG neurons cultured on the different substrates, there was no difference in the T-type channel currents (Fig. 4A-B) and volatge-dependent channel activation (Fig. 4E-F). In stark contrast, the peak amplitude of the T-type channel current was approximately three times larger in both IB4^+^ and IB4^−^ small TG neurons on the stiff substrate than on the soft substrate (Fig. 4C-D). Moreover, there was a substantial leftward shift in the voltage-dependent channel activation curve, with the V_1/2_ values decreased from −31.5 ± 3.2 mV in IB4^+^ neurons and −29.8 ± 3.2 mV in IB4^−^ neurons on the soft substrate to −49.5 ± 4.3 mV in IB4^+^ neurons and −43.9 ± 3.6 mV in IB4^−^ neurons on the stiff substrate (p < 0.05 in both cases; Fig. 4G-H). These results, taken together with no significant effect in the HVA Cav channel currents (Fig. 4D-E), indicate that stiff substrate favorably regulates the T-type channel in small TG neurons.

**Fig. 4.**
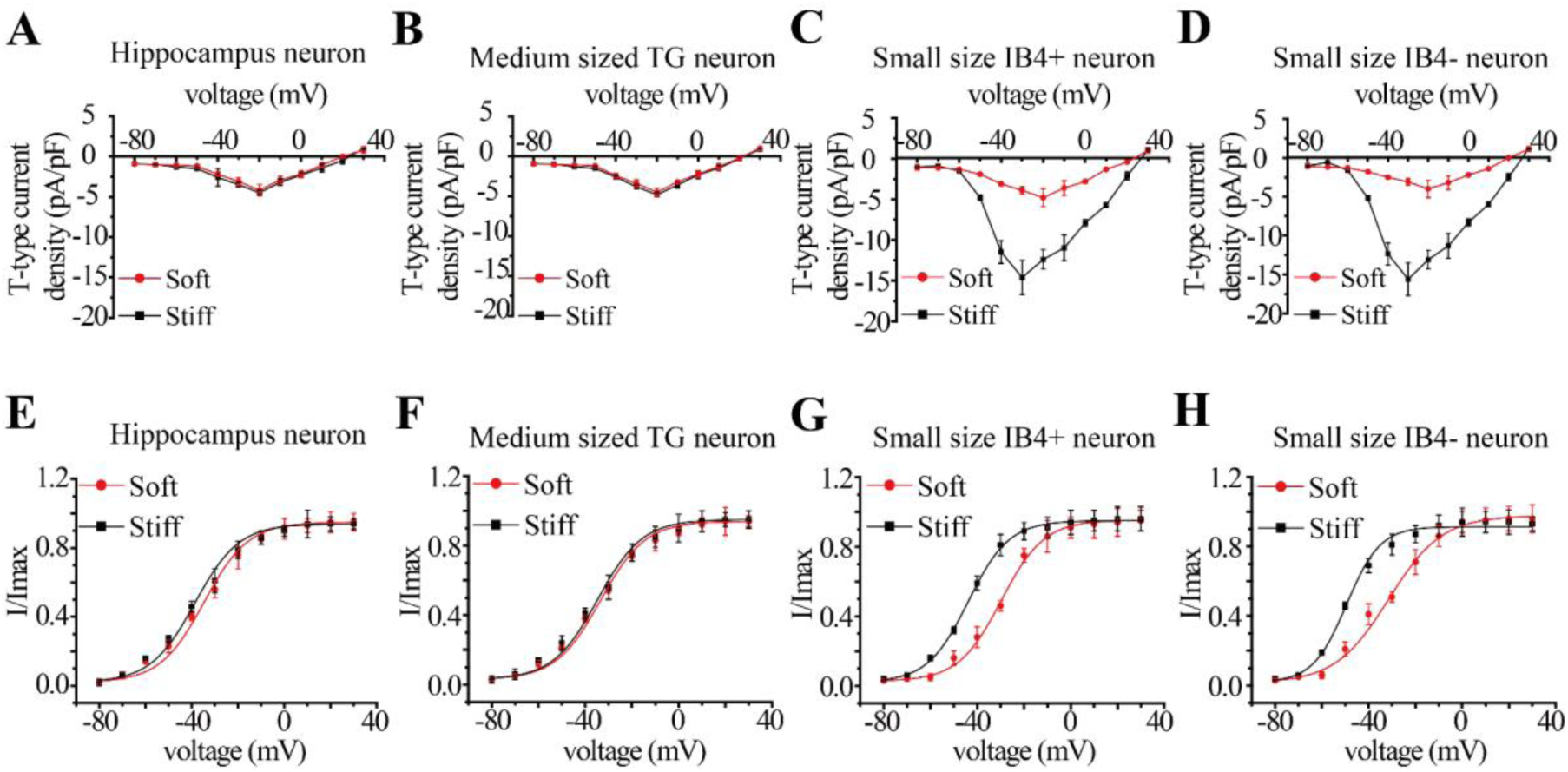
Stiff substrate up-regulates T-type channel currents in small TG neurons but not in hippocampal and medium TG neurons. **A-H.** Summary of the I-V curves of T-type channel currents (A-D) and voltage-dependent T-type channel activation determined by measuring the tail currents (E-H) from indicated neurons on the soft and stiff substrates from three preparations, with the number of neurons examined as follow: 10 hippocampal neurons (A, E), 11 medium TG neurons (B, F), 10 small IB4^+^ TG neurons (C, G) and 10 small IB4^−^ TG neurons (D, H).

### Stiff substrate enhances T-type channel-dependent excitability of small TG neurons

The T-type channel in TG neurons is well known to play a role in neuronal excitability, directly by providing a depolarizing current and/or indirectly by modulating the Ca^2+^-activated K^+^ channel current that underlies the AP repolarization, spike-frequency adaptation and after-hyperpolarization (AHP) (Zhang et al., 2002; Nelson et al., 2006). We therefore investigated whether regulation by substrate stiffness of the T-type channel currents in small TG neurons alters the neuronal excitability. As described above, there was no difference in the input resistance (R_in_) and resting membrane potential (V_rest_) for hippocampal, medium and small TG neurons cultured on the stiff and soft substrates (Tab. 2 and Tab. 3). AP-like spike responses were evoked by injection of a depolarizing current in all types of neurons (Fig. 5A-D), but stiff substrate increased the spike frequency by several folds in small TG neurons (Fig.5G-H), but not in hippocampal and medium TG (Fig.5I-J). In addition, blockage of the T-type channel with Ni^2+^ reduced the spike frequency in both IB4^+^ and IB4^−^ small TG neurons on the stiff substrate to that in neurons on the soft substrate (Fig. 5K-L). Substrate stiffness did not significantly alter the AP threshold, amplitude, width or AHP amplitude in small TG neurons (Tab. 3), suggesting that substrate stiffness has no major effect on membrane polarization or repolarization. Collectively, these results indicate that stiff substrate upregulation of the T-type channel in small TG neurons enhances the neuronal excitability.

**Fig. 5.**
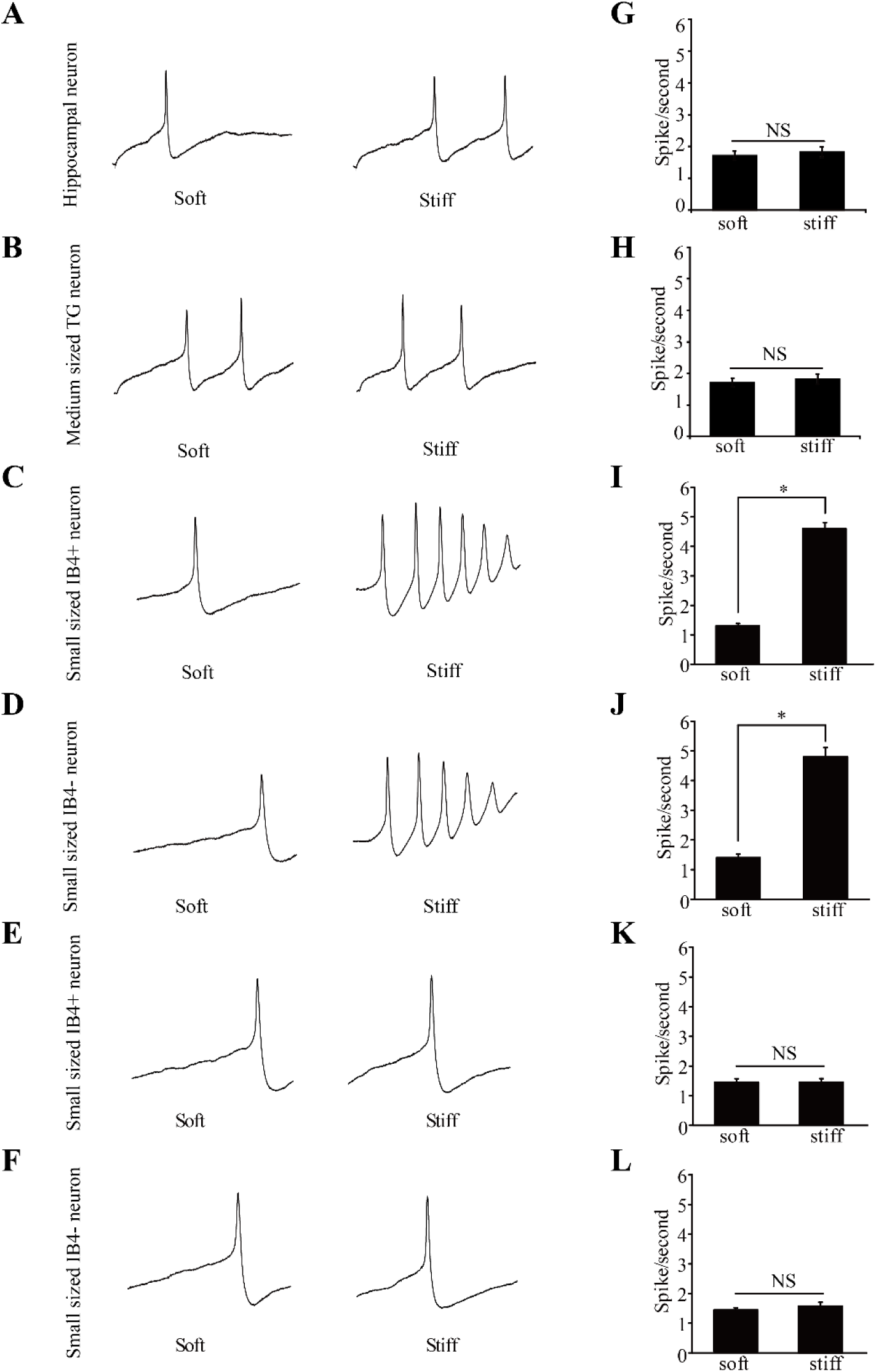
Stiff substrate increases excitability of small TG neurons via T-type channel. **A-F.** Representative recording of AP-like spikes in indicated neurons cultured on the soft and stiff substrates, evoked by injecting depolarizing currents. Note that small IB4^+^ (E) and small IB4^−^ TG neurons (F) were treated with 30 μM Ni^2+^. **G-L**. Summary of the mean spike frequency in indicated neurons on the soft and stiff substrates from three preparations as shown in A-F, with the number of neurons examined as follows: 12-14 hippocampal neurons (G), 13-14 medium TG neurons (H), 12-14 small IB4^+^ TG neurons (I), 13-14 small IB4^−^ TG neurons (J), 11-13 Ni^2+^-treated mall IB4^+^ TG neurons (K), and 10-11 Ni^2+^-treated small IB4^−^ TG neurons. *, p < 0.05, denoting significant difference between indicated groups. NS, no significant difference.

### Actomyosin mediates substrate stiffness regulation of neuronal Cav channels

As introduced above, one common mechanism by which many cells sense the mechanical properties of ECM and cell-supporting substrates is based on the traction force of actomyosin stress fibers via focal adhesions (Lee and Kumar, 2016; Kobayashi and Sokabe, 2010) and the traction force is greater in cells on the stiff substrates than that on the soft substrates (Engler et al., 2006). There is evidence to suggest that central and peripheral neurons show different sensitivity to mechanical clues (Koch et al., 2012). Here, we also revealed that substrate stiffness regulate the Cav channel types in a neuron-specific manner. Therefore, we were interested in whether both central and peripheral neurons detect substrate stiffness using actomyosin-mediated mechanosensing mechanisms. To address this, we firstly examined the effect of treatment with 3 μM blebbistatin, a low dose that causes modest inhibition of myosin II and thereby the traction force (Engler et al., 2006) on the Cav channel currents in neurons on the stiff substrate. Indeed, such treatment attenuated the total Cav channel currents in hippocampal, medium and small TG neurons on the stiff substrate and largely abolished the difference in the Cav channel current amplitude in neurons on the soft and stiff substrates (Fig. S3). Furthermore, treatment with 3 μM blebbistatin reduced the N-type channel current in hippocampal and medium TG neurons (Fig. 6A-D). In both small IB4^+^ and IB4^−^ TG neurons, such treatment also attenuated the T-type channel current (Fig. 6E-F), prevented the leftward shift in the voltage-dependent activation T-type channel curve (Fig. 6E-F), and decreased the spike frequency (Fig. 6 G-H). Overall, these results suggest a critical role for actomyosin in sensing substrate stiffness in both hippocampal and TG neurons. To further support this conclusion, we examined the effects of treatment of neurons on the soft substrate with calyculin A, which is known to induce hyperactivation and contraction of myosin (Lemmon et al., 2009; Fabian et al., 2007), and thus expected calyculin A to induce an opposite effect of blebbistatin. Treatment with 20 nM calyculin A indeed increased the Cav channel currents in hippocampal, medium and small TG neurons on the soft substrate (Fig. S4). More specifically, such treatment selectively enhanced the N-type channel current in hippocampal and medium TG neurons (Fig. 7A-D), and the T-type Ca^2+^ current and the excitability in small TG neurons (Fig. 7E-H), making the neurons on the soft substrate to be functionally similar to the neurons on the stiff substrate. Therefore, these contrasting results from treatments with blebbistatin and calyculin A consistently support critical involvement of actomyosin in both central and peripheral neurons for mechanosensing.

**Fig. 6.**
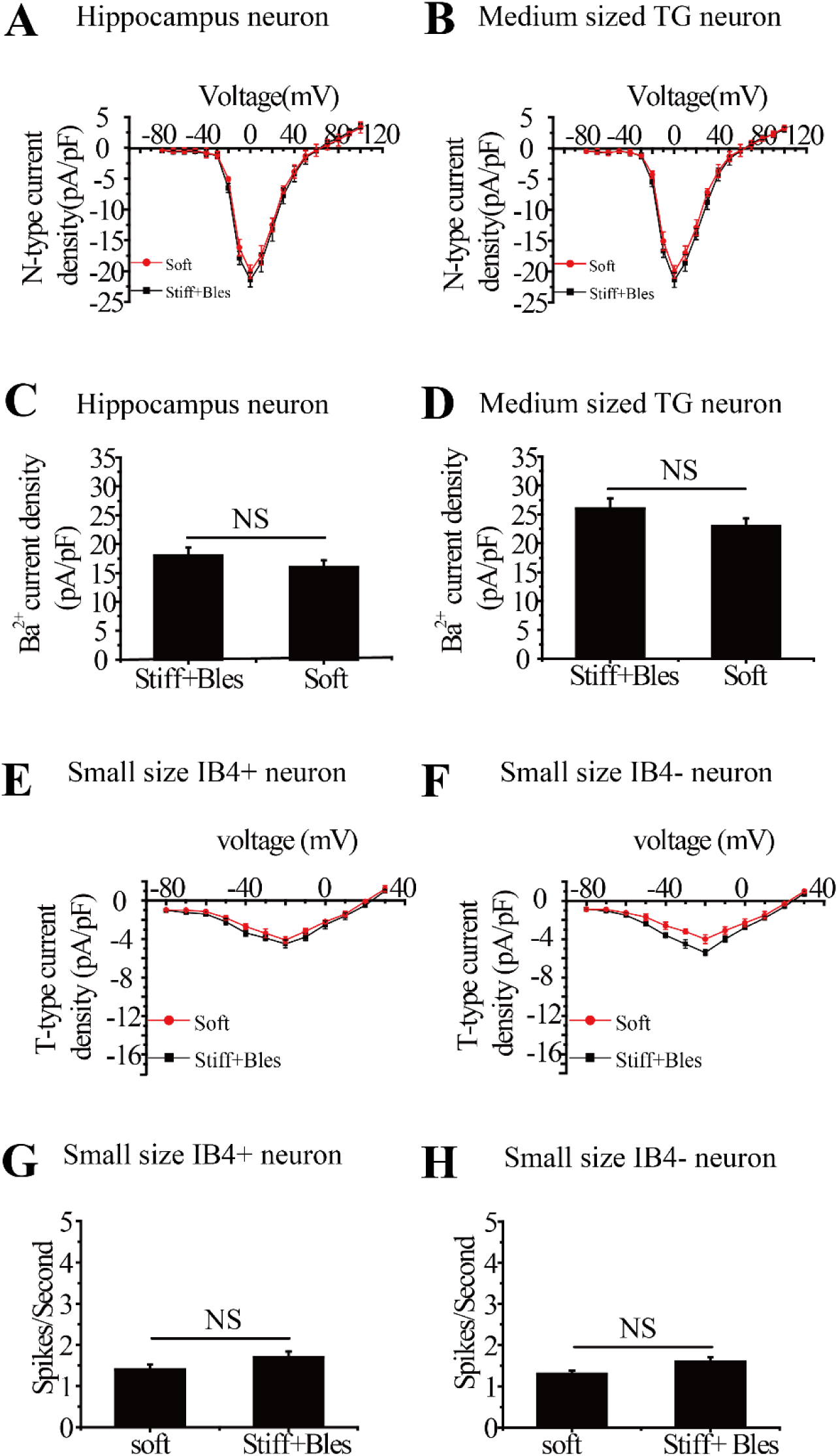
Treatment with low dose blebbistatin attenuates stiff substrate-induced upregulation of N-type channel currents in hippocampal and medium TG neurons, and T-type channel currents and excitability in small TG neurons. **A-D.** Summary of the I-V curve of N-type channel currents (A-B) and the maximal peak N-type channel currents (C-D), recorded from 11 hippocampal neurons on the stiff substrate treated with 3 μM blebbistatin and 11 hippocampal neurons on soft substrate in three preparations (A, C), and from 10 medium TG neurons on stiff substrate treated by 3 μM blebbistatin and 10 medium TG neurons on soft substrate in three preparations (B, D). **E-F**. Summary of the I-V curve of T-type Ca^2+^ channel currents, recorded from 10 small IB4^+^ TG neurons on stiff substrate treated by 3 μM blebbistatin and 10 small IB4^+^ TG neurons on soft substrate in three preparations (E), and from 11 small IB4^−^ TG neurons on stiff substrate treated by 3 μM blebbistatin and 11 small IB4^−^ TG neurons on soft substrate in three preparations (F). **G-H**. Summary of the mean spike frequency, recorded from 12 small IB4^+^ TG neurons on stiff substrate treated by 3 μM blebbistatin and 12 small IB4^+^ TG neurons on soft substrate in three preparations (G), from 12 small IB4^−^ TG neurons on stiff substrate treated by 3 μM blebbistatin and 12 small IB4^−^ TG neurons on soft substrate in three preparations (H). NS, no significant difference between indicated groups.

**Fig. 7.**
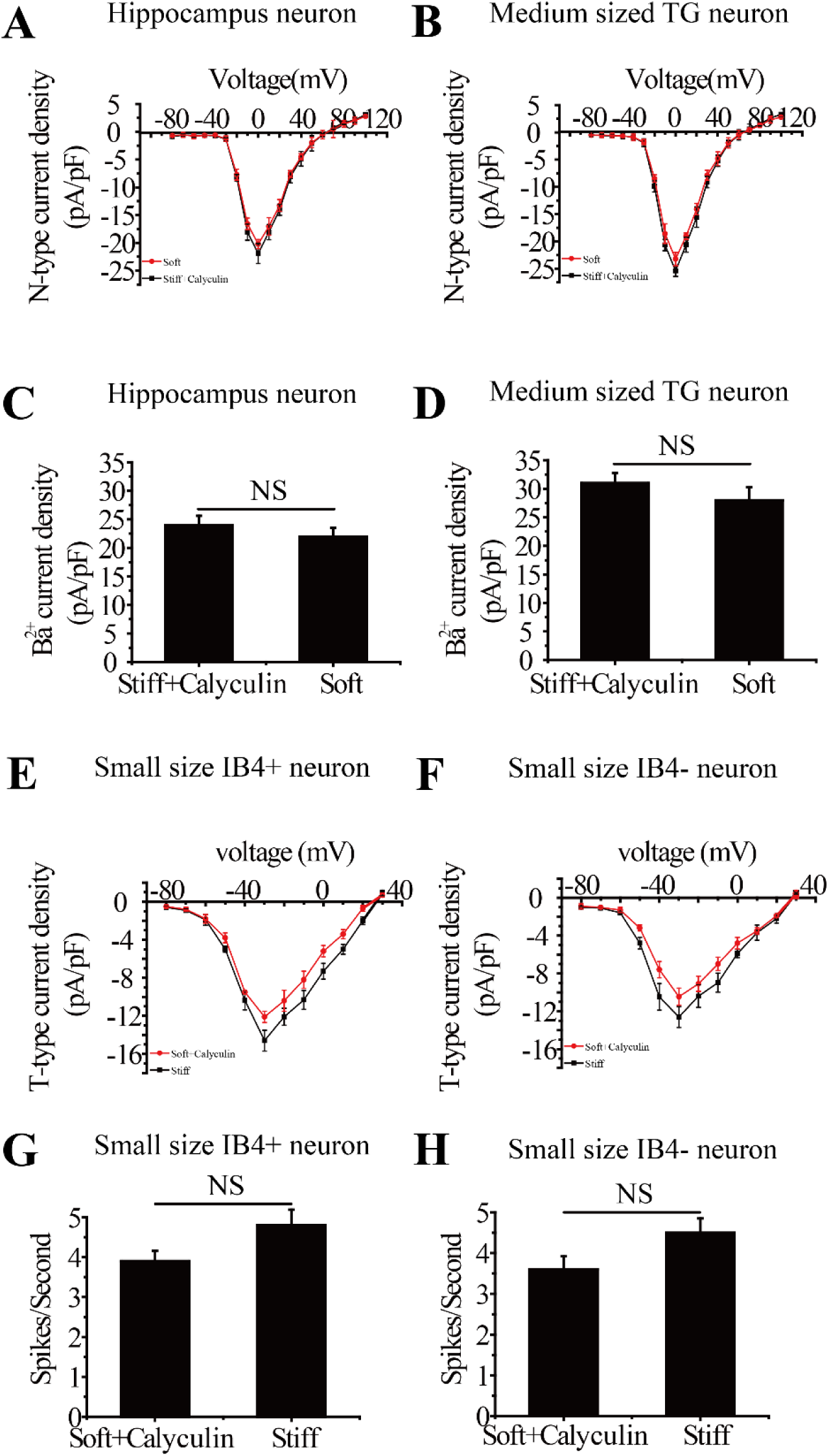
Treatment with calyculin A enhances the N-type channel currents in hippocampal and medium TG neurons, the T-type channel current and excitability in small TG neurons on the soft substrate. **A-D.** Summary of the I-V curved of N-type channel currents (A-B) and the maximal peak N-type channel currents (C-D), recorded from 10 hippocampal neurons on soft substrate treated with 20 nM calyculin A and 10 hippocampal neurons on stiff substrate in three preparations (A, C), and from 10 medium TG neurons on soft substrate treated with 20 nM calyculin A and 9 medium TG neurons on stiff substrate in three preparations (B, D). **E-F**. Summary of the I-V curve of T-type channel currents, recorded from 10 small IB4^+^ TG neurons on soft substrate treated with 20 nM calyculin A and 10 small IB4^+^ TG neurons on stiff substrate in three preparations (E), and from 11 small IB4^−^ TG neurons on the soft substrate treated with 20 nM calyculin A and 11 small IB4^−^ TG neurons on stiff substrate in three preparations (F). **G-H**. Summary of the mean spike frequency, from 12 small IB4^+^ TG neurons on soft substrate treated with 20 nM calyculin A and 12 small IB4^+^ TG neurons on stiff substrate in three preparations (G), and from 12 small IB4^−^ TG neurons on soft substrate treated with 20 nM calyculin A and 12 small IB4^−^ TG neurons on stiff substrate in three preparations (H). NS, no significant difference between indicated groups.

## Discussion

The present study provide evidence to indicate that stiff substrate preferentially regulates the N-type channel in hippocampal and medium TG neurons and the T-type channel in small TG neurons. To our best knowledge, this is the first report that mechanical regulation of the Cav channels is neuron type-specific. We also show a critical role for actomyosin-mediated mechanosensing mechanism in both central and peripheral neurons.

In this study, we have shown that the total Cav channel currents were significantly increased in hippocampal, small and medium TG neurons when cultured on the stiff substrate compared to neurons on the soft substrate (Fig. 1A-H). These findings are consistent with recent studies reported by several groups (Koch et al., 2012; Zhang et al., 2014; Lantoine et al., 2016; Bhattacharya et al., 2008; Drew et al., 2002), supporting that central and peripheral neurons can sense mechanical signals from their surroundings. Interestingly, comparison of the ratio of the Cav channel currents in neurons cultured on the soft and stiff substrates reveals significant difference between hippocampal and small TG neurons, and also between medium and small TG neurons (Fig. 1I). A clear difference between neuron types is also manifested by the finding that substrate stiffness selectively altered the voltage-dependent Cav channel activation in small TG neurons, but not in hippocampal and medium TG neurons (Fig. 2). All the Cav channel types are functionally expressed in all neuron types examined in this study (Fig.3 and Fig. 4). These differences may arise from different mechanosensing mechanisms, different Cav channel types under the regulation, or both. Our results provide strong evidence that substrate stiffness preferentially regulates the N-type channel in hippocampal and medium TG neurons (Fig. 3) and, by contrast, the T-type channel in small TG neurons (Fig. 4). Substrate stiffness regulation of different types of Cav channels in hippocampal and small TG neurons seems to be well aligned with the key role of these two channels in the function of these neuron types. The N-type channel provides the major Ca^2+^ source for neurotransmission, particularly in the central nervous system (Cao et al., 2004; Chaplan et al., 1994; Rycroft et al., 2007). Consistently, stiff substrate has been recently shown, in experiments on hippocampal (Zhang et al., 2014) and cortical neurons (Lantoine et al., 2016), to enhance neuronal network formation and activity. On the other hand, the T-type channel readily opens in resposne to brief and modest membrane depolarization and is critically involved in regulating the excitability of sensory neurons (Tao et al., 2012; Fang et al., 2007; Bourinet et al., 2005). In agreement with these notions, we showed in this study that stiff substrate increased the excitability of small TG neurons in a T-type channel-dependent manner (Fig. 5 K-L). Recent studies have reported different mechanosensitivity of central and peripheral neurons in that DRG neurite displayed maximal outgrowth on the soft substrate, whereas hippocampal neurite outgrowth is insensitive to substrate stiffness (Koch et al., 2012; Koser et al., 2016). Like TG neurons, DRG neurons are heterogeneous in size and associated with distinctive functional roles. There is evidence that medium TG neurons are more sensitive to stretch intensity than small TG neurons (Rosoff et al., 2004; Bhattacharya et al., 2008). Consistently with difference in mechanosensitivity, our results show that the Cav channel current ratio in medium TG on the stiff versus soft substrates was significantly greater than that in small TG medium (Fig. 1I). Furthermore, we have revealed that substrate stiffness regulation targets the N-type, not the T-type channel in medium TG neurons and has no effect on the neuronal excitability (Fig. 3 and Fig. 4). A previous report suggested that small IB4^+^ TG neurons were less sensitive to externally applied mechanical stimuli than IB4^−^ neurons (Drew et al., 2002; Bhattacharya et al., 2008). In this present study we found no difference in the regulation by substrate stiffness of small IB4^+^ and IB4^−^ TG neurons (Figs.1-5). Overall, the present study provides evidence to suggest neuron type-specific regulation of Cav channels in the central and peripheral neurons. Further investigations are warranted to provide a clear understanding of such differential mechanical regulation of Cav channels and neuronal excitability. It is worth pointing out that hippocampal neuronal preparations often contained glial cells, which are known to have bidirectional communications with neurons (Lee and Haydon, 2011; Halassa et al., 2007), and we cannot exclude the possibility that glial cells may confer hippocampal neurons with an additional and indirect mechanosensing capacity.

The actomyosin-mediated traction force mechanism is most commonly used among many other known mechanosensing mechanisms (Lee and Kumar, 2016; Kobayashi and Sokabe, 2010; Engler et al., 2006). In this study, we examined involvement of the actomyosin-mediated mechanosensing mechanism in the differential regulation of Cav channels in the central and peripheral neurons by substrate stiffness, using blebbistatin and calyculin A to inhibit or enhance myosin function. Treatment of neurons cultured on the stiff substrate with low dose blebbistatin lowered the N-type channel currents in hippocampal and medium TG neurons, and also the T-type channel currents and T-type channel-dependent excitability in small TG neurons to the same levels in neurons on the soft substrate (Fig. 6). By contrast, treatment of neurons cultured on the soft substrate with calyculin A increased the N-type channel currents in hippocampal and medium TG neurons, and also the T-type channel currents and T-type channel-dependent excitability in small TG neurons to the same levels in neurons on the stiff substrate (Fig. 7). Such contrasting results are highly consistent in supporting critical involvement of actomyosin as part of the mechanisms for mechanical regulation of different Cav channels and associated neuronal functions in both central and peripheral neurons.

In summary, this study reports the first evidence to show neuron type-specific regulation of different Cav channels in the central and peripheral neurons by substrate stiffness. While the stiffness of the substrates used is relatively high, whoever, recent report shown that neurons sense stiffness gradients and the sensing was independent of the absolute stiffness (Koser et al., 2016). Thus, the findings reported in this study provide very useful information for development of novel interfaces for neural tissue engineering and regeneration.

## Author contributions

H.C.Z. and L.-H.J. conceived the research, designed the experiments and prepared the manuscript. S.S.L., Y.Y. X.A.W, Y.C.Q and L.Y.H performed the experiments. Y.Y. and X.A.W. analyzed the data. All authors discussed the results and commented on the manuscript.

## Acknowledgments

The work was support by research grants (No.11072132, 11472159 and No.11272184) from National Natural Science Foundation of China (to H.C.Z.) and Xinxiang Medical University, Department of Education, and Office of Foreign Experts Affairs, Henan Province, China (to L.-H.J.).

## Additional information

Competing financial interests: the authors declare no competing financial interests.

## Supplementary data

**Fig. S1.**
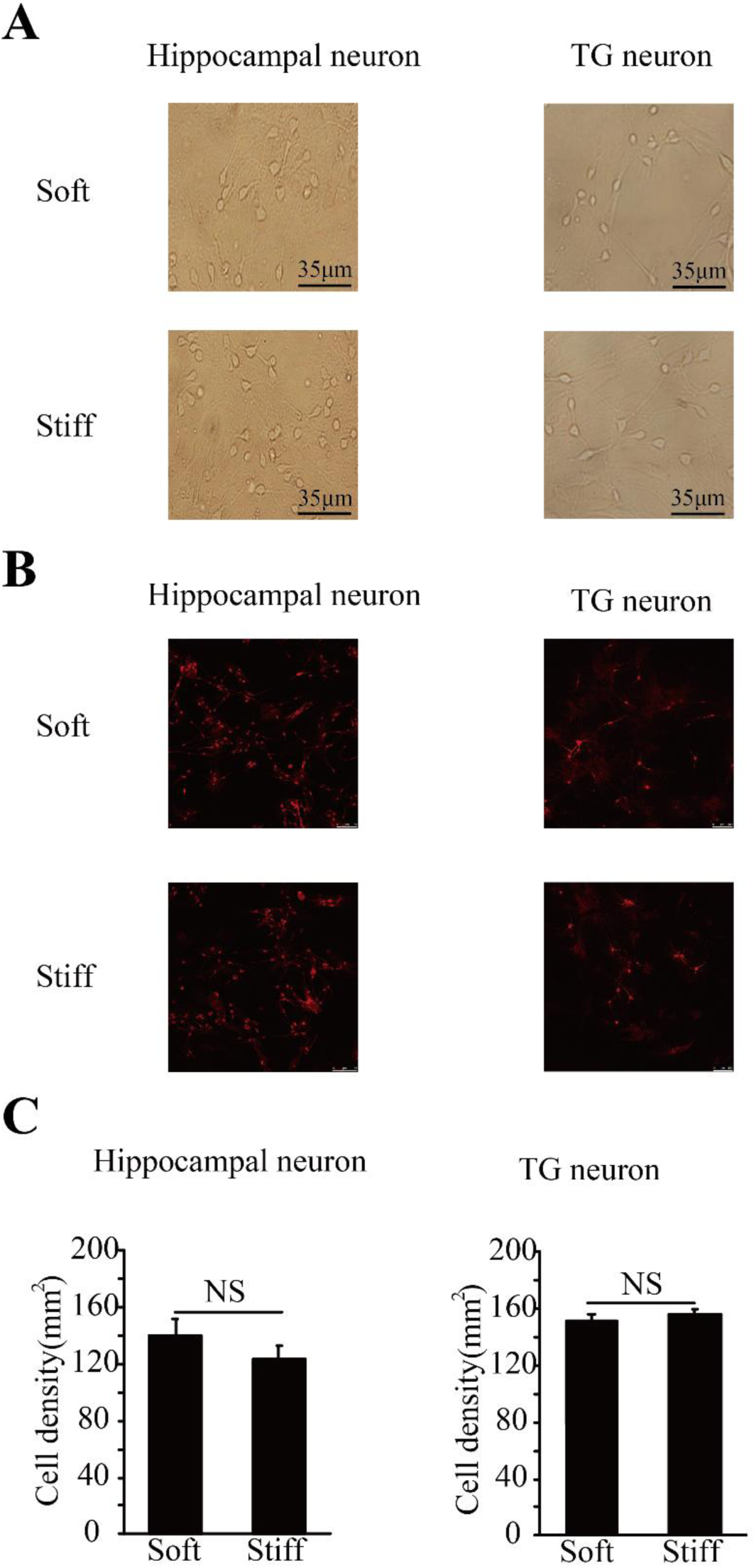
No effect of substrate stiffness on neuronal cell density in hippocampal and TG neuronal cultures. (**A**) Representative optical microscopic photos of hippocampal and TG neurons. (**B**) Fluorescent confocal images showing MAP2 staining of hippocampal and TG neurons. (**C**) Summary of neurons cell density determined from images, as shown in A, from 10 independent preparations for each case. NS, no significant difference.

**Fig. S2.**
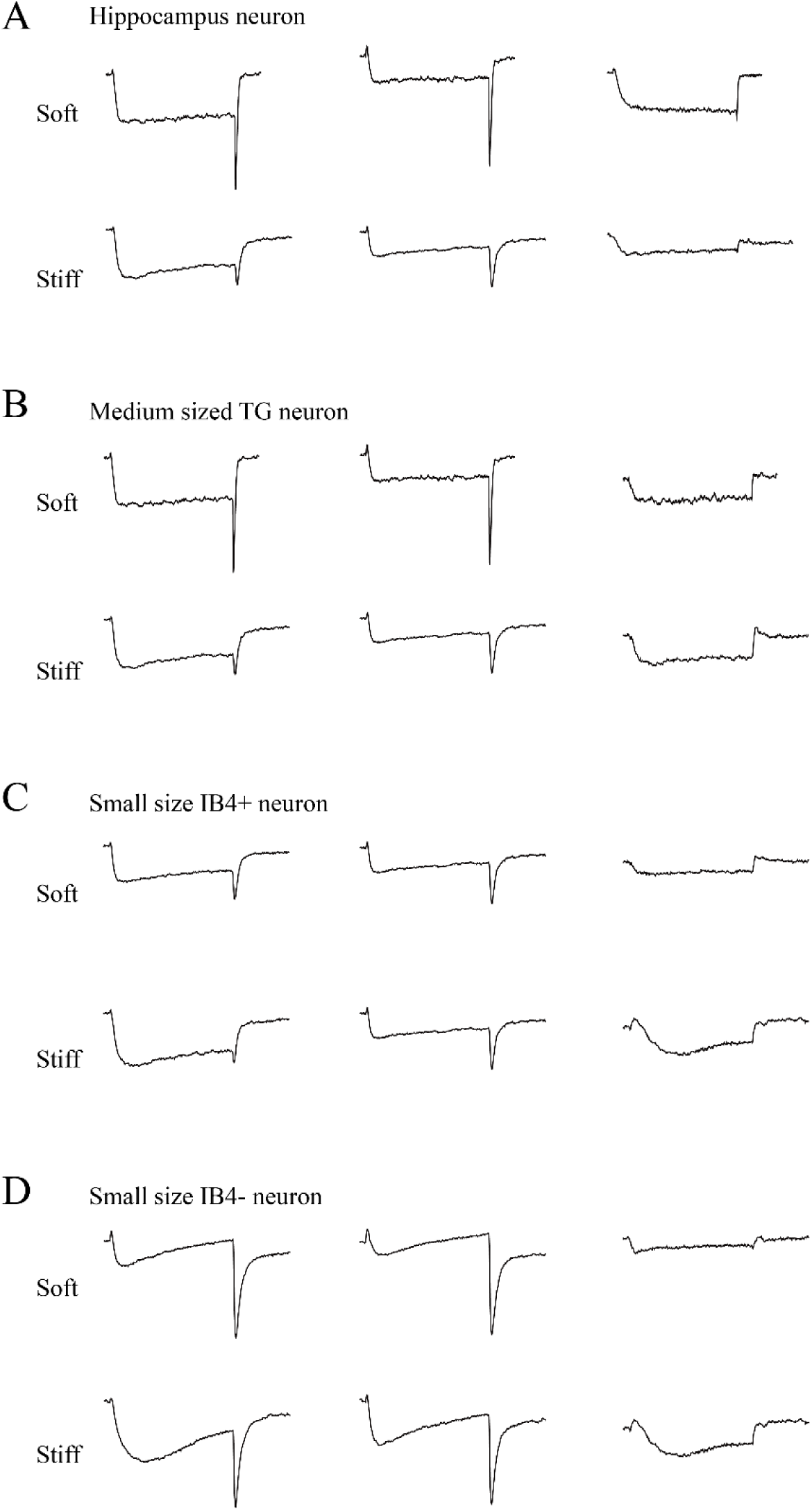
Isolation of T-type channel currents in hippocampal neurons and TG neurons on soft and stiff substrates. Representative recordings from hippocampal neurons (A), and medium sized TG neurons (B), small IB4^+^ TG neurons (C), and IB4^−^ TG neurons (D) cultured on soft and stiff substrates in the presence of HVA channel inhibitors (0.5 μM ω-Aga-IVA, 2 μM ω-conotoxin-GVIA, 10 μM nifedipine and 100 μM Cd^2+^). *Left:* Ba^2+^ currents elicited by depolarization from the V_h_ of −110 mV to −40 mV. *Middle:* Ba^2+^ currents elicited by depolarization from the V_h_ of −60 mV to −40 mV. *Right:* LVA T-type channel currents obtained by subtracting the current measured from the V_h_ of −60 mV (middle) from matched current measured from the V_h_ of −110 mV (left).

**Fig. S3.**
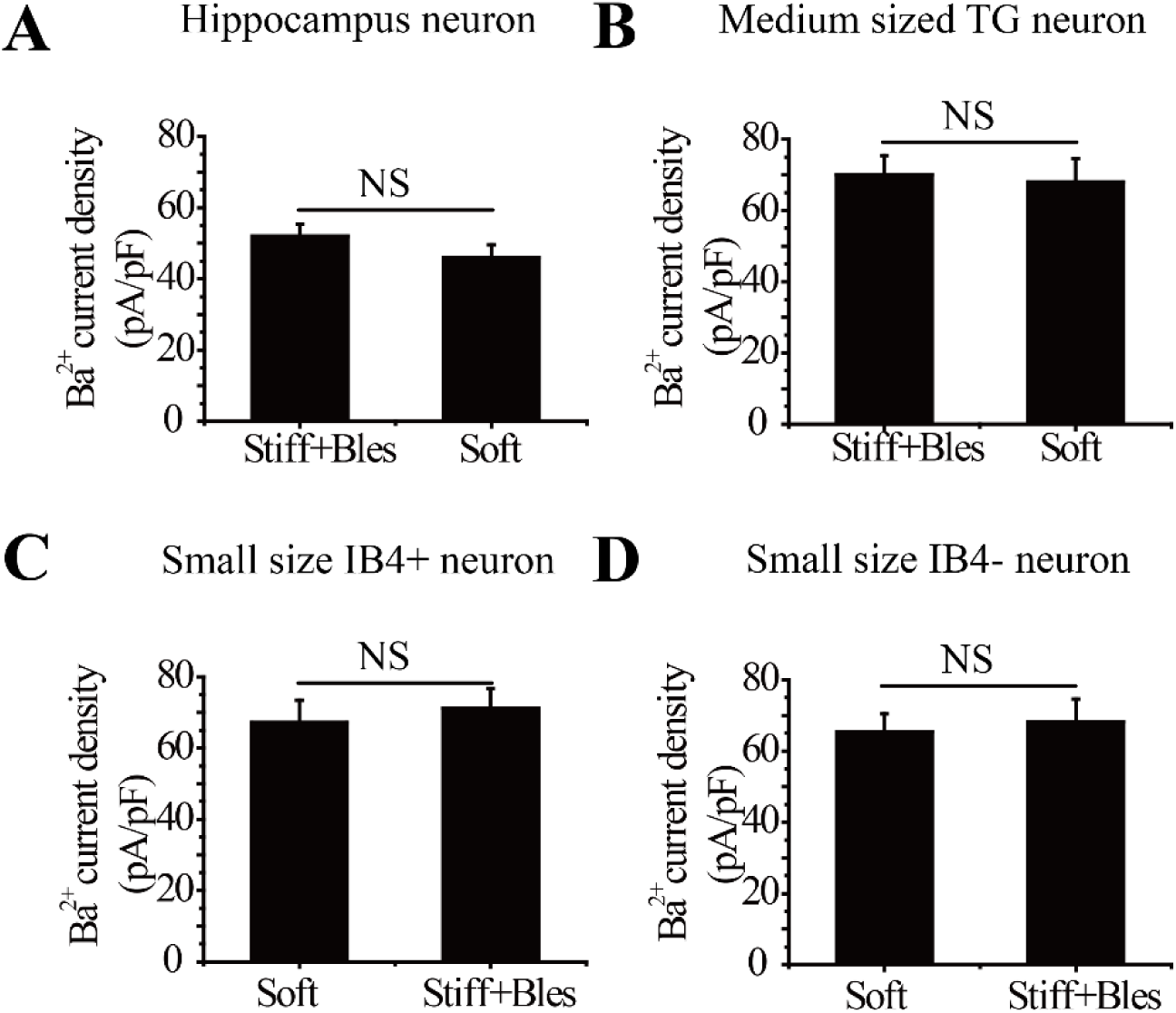
Comparison of Cav channel current density in hippocampal and TG neurons on the stiff substrate treated with low dose blebbistatin and in neurons on the soft substrate. Summary of the peak Cav channel current density, from 11 hippocampal neurons on stiff substrate treated with 3 μM blebbistatin and 11 hippocampal neurons on soft substrate in three preparations (A), from 10 medium neurons on stiff substrate treated with 3 μM blebbistatin and 10 medium TG neurons on soft substrate in three preparations (B), from 10 small IB4^+^ TG neurons on stiff substrate treated with 3 μM blebbistatin and 10 small IB4^+^ TG neurons on soft substrate in three preparations (C), and from 11 small IB4^−^ TG neurons on stiff substrate treated with 3 μM blebbistatin and 11 small IB4^−^ TG neurons on soft substrate in three preparations. NS, no significant difference.

**Fig. S4.**
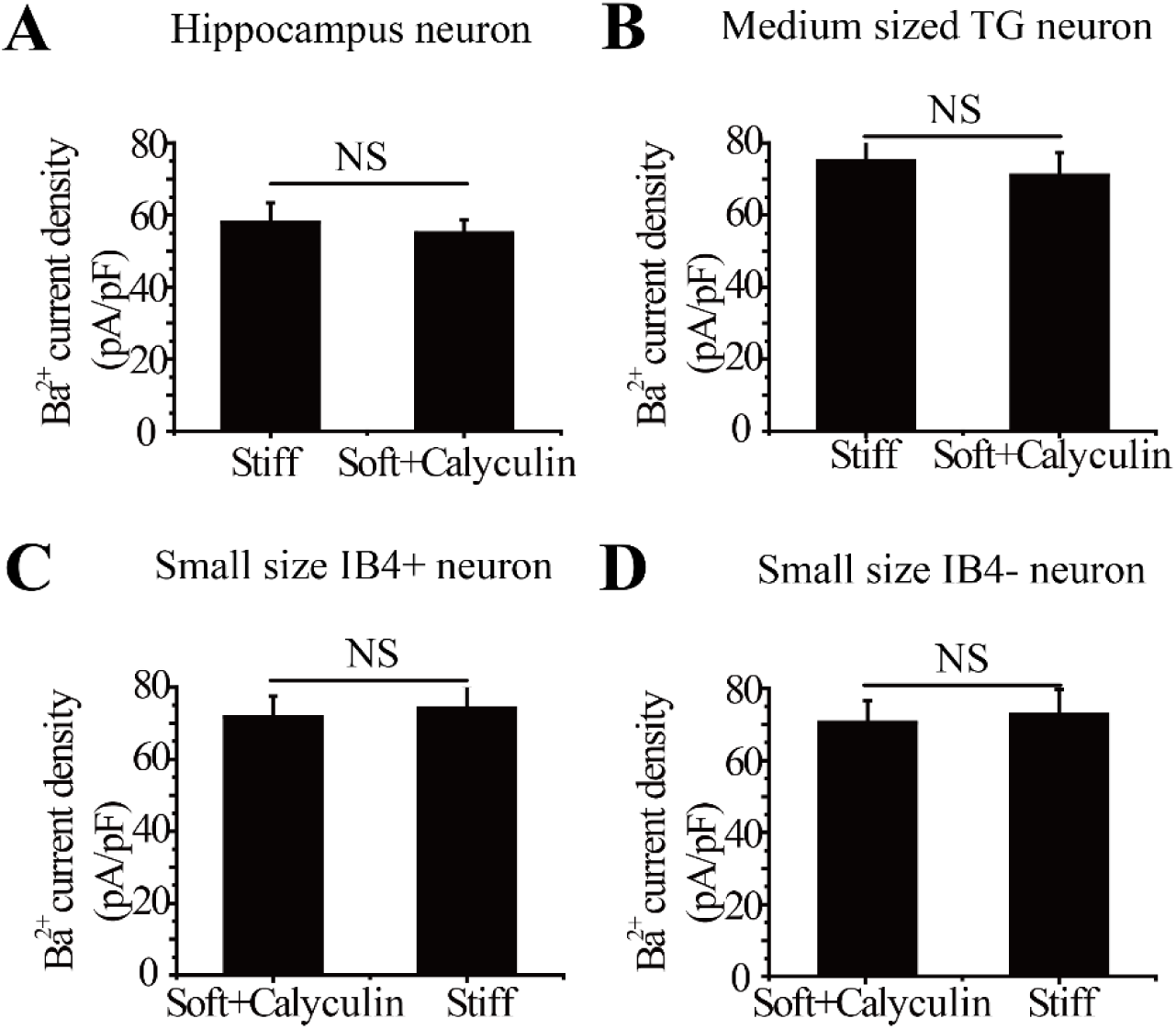
Comparison of Cav channel current density in hippocampal and TG neurons on the soft substrate treated with calyculin A and in neurons on the stiff substrate. Summary of the peak Cav current density, from 11 hippocampal neurons on soft substrate treated with 20 nM calyculin A and 11 hippocampal neurons on stiff substrate in three preparations (A), from 11 medium neurons on soft substrate treated with 20 nM calyculin A and 11 medium TG neurons on stiff substrate in three preparations (B), from 10 small IB4^+^ TG neurons on soft substrate treated with 20 nM calyculin A and 10 small IB4^+^ TG neurons on stiff substrate in three preparations (C), and from 11 small IB4^−^ TG neurons on soft substrate treated with 20 nM calyculin A and 11 small IB4^−^ TG neurons on stiff substrate in three preparations. NS, no significant difference.

## References

Beaudoin, G., Lee, S.H., Singh, D., Yuan, Y., Ng, Y.G., Reichardt, L.F. and Arikkath, J. (2012). Culturing pyramidal neurons from the early postnatal mouse hippocampus and cortex. Nat. Protoc. 7, 1741–1754.

Bhattacharya, M.R.C., Bautistab, D.M., Wu, K., Haeberle, H., Lumpkin, E.A. and Julius, D. (2008). Radial stretch reveals distinct populations of mechanosensitive mammalian somatosensory neurons. Proc. Natl. Acad. Sci. USA. 105, 20015–20020.

Bourinet, E., Alloui, A., Monteil, A., Barrere, C., Couette, B., Poirit, O., McRory, J., Snutch, T.P., Eschalier, A. and Nargeot, J. (2005). Silencing of the Cav3.2 T-type calcium channel gene in sensory neurons demonstrates its major role in nociception. EMBO. J. 24, 315–324.

Cao, Y.Q., Piedras-Renteria, E.S., Smith, G.B., Chen, G., Harata, N.C. and Tsien, R.W. (2004). Presynaptic Ca^2+^ channels compete for channel type-preferring slots in altered neurotransmission arising from Ca^2+^ channelopathy. Neuron. 43, 387–400.

Catterall, W.A. and Swanson, T.M. (2015). Structural basis for pharmacology of voltage-gated sodium and calcium channels. Mol. Pharmacol. 88, 141–150.

Cellot, G., Toma, F.M., Varley, Z.K., Laishram, J., Villari, A., Quintana, M., Cipollone, S., Prato, M. and Ballerini, L. (2011). Carbon nanotube scaffolds tune synaptic strength in cultured neural circuits: novel frontiers in nanomaterial-tissue interactions. J. Neurosci. 31, 12945–12953.

Chaplan, S.R., Pogrel, J.W. and Yaksh, T.L. (1994). Role of voltage-dependent calcium channel subtypes in experimental tactile allodynia. J. Pharmacol. Exp. Ther. 269,1117–1123.

Cheong, H. and Shin, H.S. (2013). T-type Ca^2+^ channels in normal and abnormal brain functions. Physiol. Rev. 93, 961–992.

Drew, L.J., Wood, J.N. and Cesare, P. (2002). Distinct mechanosensitive properties of capsaicin-sensitive and insensitive sensory neurons. J. Neurosci. 22, RC228.

Engler, A.J., Sen, S., Sweeney, H.L. and Discher, D.E. (2006). Matrix elasticity directs stem cell lineage specification. Cell. 126, 677–689.

Fang, Z., Park, C.K., Li, H.Y., Kim, H.Y., Park, S.H., Jung, S.J., Kim, J.S., Monteil, A., Oh, S.B. and Miller, R.J. (2007). Molecular basis of Ca(v)2.3 calcium channels in rat nociceptive neurons. J. Biol. Chem. 282, 4757–4764.

Franze, K and Guck, J. (2010). The biophysics of neuronal growth. Rep. Prog. Phys. 73, 094601.

Fabbro, A., Bosi, S., Ballerini, L. and Prato, M. (2012). Carbon nanotubes: artificial nanomaterials to engineer single neurons and neuronal networks. ACS. Chem. Neurosci. 15, 611–618.

Fabian, L., Troscianczuk, J. and Forer, A. (2007). Calyculin A, an enhancer of myosin, speeds up anaphase chromosome movement. Cell & Chromosome. 6, 1.

Georges, P.C., Miller, W.J., Meaney, D.F., Sawyer, E.S. and Janmey, P.A. (2006). Matrices with compliance comparable to that of brain tissue select neuronal over glial growth in mixed cortical cultures. Biophys. J. 90, 3012–3018.

Halassa, M.M., Fellin, T. and Haydon, P.G. (2007). The tripartite synapse: roles for gliotransmission in health and disease. Trends. Mol. Med. 13, 54–63.

Kobayashi, T. and Sokabe, M. (2010). Sensing substrate rigidity by mechanosensitive ion channels with stress fibers and focal adhesions. Curr. Opin. Cell. Biol. 22, 669–676.

Koch, D., Rosoff, W.J., Jiang, J.J., Geller, H.M. and Urbach, J.S. (2012). Strength in the periphery: growth cone biomechanics and substrate rigidity response in peripheral and central nervous system neurons. Biophys. J. 102, 452–460.

Koser, D.E., Thompson, A.J., Foster, S.K., Dwivedy, A., Pillai, E.K., Sheridan, G.K., Svoboda, H., Viana, M., Costa, L.D., Guck, J., Holt, C.E. and Franze, K. (2016). Mechanosensing is critical for axon growth in the developing brain. Nat. Neurosci. 19, 1592–1598.

Kostic, A., Sap, J. and Sheetz, M.P. (2007). RPTPα is required for rigidity-dependent inhibition of extension and differentiation of hippocampal neurons. J. Cell. Sci. 120, 3895–3904.

Lantoine, J., Grevesse, T., Villers, A., Delhaye, G., Mestdagh, C., Versaeve, M., Mohammed, D., Bruyere, C., Alaimo, L. and Lacour, S.P. (2016), Matrix stiffness modulates formation and activity of neuronal networks of controlled architectures. Biomaterials. 89, 14–24.

Lee, S. and Kumar, S. (2016). Actomyosin stress fiber mechanosensing in 2D and 3D. F1000Research. 5, 10.12688/f1000research.8800.1.

Lee, S.Y. and Haydon, P.G. (2011). A Cytokine-dependent switch for glial-neuron interactions. Neuron. 69, 835–837.

Lemmon, C.A., Chen, C.S. and Romer, L.H. (2009) Cell traction forces direct fibronectin matrix assembly. Biophys. J. 96, 729–738.

Lu, Y.B., Franze, K., Seifert, G., Steinhauser, C., Kirchhoff, F., Wolburg, H., Guck, J., Janmey, P., Wei, E.Q., Kas, J. and Reichenbach, A. (2006). Viscoelastic properties of individual glial cells and neurons in the CNS. Proc. Natl. Acad. Sci. USA. 103, 17759–17764.

Nelson, M.T., Todorovic, S.M. and Perez-Reyes, E. (2006). The role of T-type calcium channels in epilepsy and pain. Curr. Pharm. Des. 12, 2189–2197.

Previtera, M.L., Hui, M., Verma, D., Shahin, A.J., Schloss, R. and Langrana, N.A. (2013). The effects of substrate elastic modulus on neural precursor cell behavior. Ann. Biomed. Eng. 41, 1193–1207.

Previtera, M.L., Langhammer, C.G., Langrana, N.A. and Firestein, B.L. (2010). Regulation of dendrite arborization by substrate stiffness is mediated by glutamate receptors. Ann. Biomed. Eng. 38, 3733–3743.

Proft, J. and Weiss, N. (2015). G Protein regulation of neuronal calcium channels: Back to the future. Mol. Pharmacol. 87, 890–906.

Rosoff, W. J., Urbach, J.S. and Goodhill, G.J. (2004) A new chemotaxis assay shows the extreme sensitivity of axons to molecular gradients. Nat. Neurosci. 7, 678–682.

Rycroft, B.K., Vikman, K.S. and Christie, M.J. (2007). Inflammation reduces the contribution of N-type calcium channels to primary afferent synaptic transmission onto NK1 receptor-positive lamina I neurons in the rat dorsal horn. J. Physiol. 580, 883–894.

Schiffhauer, E.S., Luo, T.Z., Mohan, K., Srivastava, V., Qian, X.Y., Griffis, E.R., Iglesias, P.A. and Robinson, D.N. (2016). Mechanoaccumulative Elements of the Mammalian Actin Cytoskeleton. Curr. Biol. 26, 1473–1479.

Sur, S., Newcomb, C.J., Webber, M.J. and Stupp, S.I. (2013). Tuning supramolecular mechanics to guide neuron development. Biomaterials. 34, 4749–4757.

Tang, M.L., Song, Q., Li, N., Jiang, Z.Y., Huang, R. and Cheng, G.S. (2013). Enhancement of electrical signaling in neural networks on graphene films. Biomaterials. 34, 6402–6411.

Tao, J., Liu, P., Xiao, Z.M., Zhao, H.C., Gerber, B.R. and Cao, Y.Q. (2012). Effects of familial hemiplegic migraine type 1 mutation T666M on voltage-gated calcium channel activities in trigeminal ganglion neurons. J. Neurophysiol. 107, 1666–1680.

Zamponi, G.W. (2016). Targeting voltage-gated calcium channels in neurological and psychiatric diseases. Nat. Rev. Drug. Discov. 15, 19–34.

Zhang, M.G., Cao, Y.P., Li, G.Y. and Feng, X.Q. (2014). Spherical indentation method for determining the constitutive parameters of hyperelastic soft materials. Biomech. Model. Mechanobiol. 13: 1–11.

Zhang, Q.Y., Zhang, Y.Y., Xie, J., Li, C.X., Chen, W.Y., Liu, B.L., Wu, X.A., Li, S.N., Huo, B., Jiang, L.H. and Zhao, H.C. (2014). Stiff substrates enhance cultured neuronal network activity. Sci. Rep. 4, 10.1038/srep06215.

Zhang, Y., Mori, M., Burgess, D.L. and Noebels, J.L. (2002). Mutations in high-voltage-activated calcium channel genes stimulate low-voltage-activated currents in mouse thalamic relay neurons. J. Neurosci. 22, 6362–6371.

